# Age dependent trans-cellular propagation of human tau aggregates in *Drosophila* disease models

**DOI:** 10.1101/2020.03.28.013136

**Authors:** Aqsa, Surajit Sarkar

## Abstract

Tauopathies is a class of neurodegenerative disorders which involves the transformation of physiological tau into pathogenic tau. One of the prime causes reported to drive this conversion is tau hyperphosphorylation and the subsequent propagation of pathogenic protein aggregates across the nervous system. Although past attempts have been made to deduce the details of tau propagation, yet not much is known about its mechanism. A better understanding of this aspect of disease pathology can prove to be beneficial for the development of diagnostic and therapeutic approaches. In our work, we utilize the plethora of advantages procured by *Drosophila* to introduce a novel *in-vivo* tauopathy propagation model. For the first time, we demonstrate that the human tau (h-tau) possesses an intrinsic property to spread trans-cellularly in the fly nervous system irrespective of the tau allele or the neuronal tissue type. Aggregate migration restricted by targeted down-regulation of a specific kinase, elucidates the role of hyper-phosphorylation in its movement. On the contrary to the previous models, the present system enables a rapid, convenient and robust *in-vivo* study of tau migration pathology. Henceforth, the developed model would not only be immensely helpful in uncovering the mechanistic in-depths of tau propagation pathology but also aid in modifier and/or drug screening for amelioration of tauopathies.

## Introduction

Neurodegenerative disorders encompass a class of disorders termed as “Tauopathies”, which being heterogeneous in nature, itself encloses a plethora of diseases characterized by the accumulation of aggregated tau protein [1]. Tau in its physiological form has many roles to play among which the most crucial is the microtubule binding and its stabilization, and axonal transport. Its conversion into pathogenic form, leads to its aggregation and formation of neurofibrillary tangles (NFTs) which eventually form the basis of the tau pathogenesis [2]. Tau aggregation is driven by a variety of factors including post translation modifications such as phosphorylation, mutations of the *MAPT* (microtubule associated protein tau) gene, injury of the brain, atypical expression of tau isoforms, and spread from the neighbouring cells [3].

Among the above triggers, the spreading of tau from neighbouring cells is an idea that developed since staging of Alzheimer’s pathology was studied [4, 5]. The observation that tau progresses throughout the brain *via* synaptically connected regions led to the hypothesis that tau propagates and spreads via neuronal circuits [5]. The propagation of tau has been distinguished into three major stages with the intracellular tau; (i) being secreted into extracellular space, followed by its (ii) uptake by recipient neurons and finally (iii) formation of novel intracellular aggregates in the recipient cells [6]. Further evidences over the years have strengthened this hypothesis with demonstrations like, injection of brain extracts of mutant P301S mice expressing wild type human tau (exhibit tau pathology) into the hippocampus of transgenic wild type tau-expressing mice (that do not manifest tau pathology) led to development and spread of tau filamentous pathology in it [7]. It has also been demonstrated that insoluble misfolded human tau is capable of inducing and propagating neurofibrillary pathology in rat model expressing non-mutated human truncated tau protein [8]. Several studies have suggested that tau entities do progress in a “prion like” fashion [5, 6, 9]. Although, immense amount of work has been contributed to the understanding of seed and spread mechanism of tau pathology via various model systems [8, 10, 11], yet it is still a field of potential exploration. Therefore, present hour demands the creation of such *in vivo* model systems which can facilitate the in-depth study of the tau propagation with ease and reproducibility.

Hence, to gain a better perspective of the mechanism of tau propagation and its role in disease pathology, we developed a novel *Drosophila in-vivo* tauopathy model. *Drosophila* disease models have been widely accepted as they recapitulate many structural and functional aspects of human disease pathology and offer innumerable advantages over other model systems [12]. In our study, using the *Drosophila in-vivo* model, we demonstrate that human tau propagates trans-cellularly from the site of targeted expression to the additional regions of the adult fly brain by the factor of age. Hence, our finding is in line with the previous reports which also proclaim the migration of tau pathology in other model systems [13, 14]. Interestingly, no propagation was noted at the larval stage in spite of a robust expression of tau. We further articulate that enhanced tau instability and eventually aggregation due to increased tau phosphorylation over age, is a driving force that over time leads to its propagation. Further, rescue in the extent of propagated tau caused by the down regulation of its phosphorylation through the introduction of a genetic modifier, validates our hypothesis, that hyper-phosphorylation does aid tau propagation. Henceforth, the *Drosophila* model thus created provides a convenient and coherent system to study the detailed mechanism of tau spreading and can be explored further, thereby aiding in the diagnostic and therapeutic fields of research.

## Materials and methods

### Fly stocks

The *Drosophila* transgenic stocks utilised were propagated on media composed of corn/agar/yeast at 24 ± 1 °C under a 12h light/dark cycle. The transgenic stocks of *Drosophila* consisting of human tau (0N/4R, four microtubule binding domains); *UAS-tau*^*WT*^ (wild type form of tau) and *UAS-tau*^*v337*^ (Val^337^→Met) were used [15]. The others utilized were *UAS-GFP* (Bloomington stock no. 1521), *GMR-Gal4* [16], *w*; P{ple-GAL4.F}3* (Bloomington stock no. 8848) and *UAS-GSK3β RNAi* (VDRC stock no. 101538).

### Histology and immunohistochemistry

For immunohistochemical staining, fixation of the decapitated heads was performed in 4% paraformaldehyde (PFA), followed by dehydration in increasing concentrations of 50% to 100% of ethanol, and embedding in paraffin wax (Sigma Aldrich, USA). The serialized frontal head sections of 15 µm thickness were obtained using a microtome (Shandon™ Finesse ™ 325 Manual Microtome, Thermo Scientific, USA) on slides coated with 0.1% poly-L-Lysine (Sigma-Aldrich, USA). Followed by dewaxing using xylene and rehydration, tissue sections were blocked at room temperature for 2 hours. Primary antibody incubation was performed at 4 °C for overnight, followed by 1X PBST washes and secondary antibody incubation at room temperature. Counterstaining and subsequent mounting was performed in DAPI (5μg/ml, Roche Diagnostics GmbH, Germany) and Prolong Gold antifade reagent (Thermofisher Scientific, US) respectively.

For immunostaining of third instar larval eye discs with brain, following dissection, tissues were fixed in 4% PFA followed by 1X PBST washes, overnight primary antibody incubation and subsequent steps were identical to the above protocol.

For co-immunostaining in the whole mount brain and sections, the protocol was followed till 1X PBST washes on the consecutive day of the first primary antibody incubation. Eventually, tissues were again incubated in the second primary antibody for consecutive overnight. 1X PBST washes on the following day were preceded by secondary antibody incubations at room temperature identical to the sequence of the primary antibody incubations. Finally, the tissues were counter stained in DAPI (5μg/ml, Roche Diagnostics GmbH, Germany) and mounted in Prolong Gold antifade reagent (Thermofisher Scientific, US).

The primary antibodies utilised were anti-human tau (5A6, 1:1000, DSHB, USA), anti-TH (1:1000, Immunostar, USA) and anti-tau (1:200, Booster, USA), anti-GFP (1:500, Booster, USA), anti-human PHF-tau^Thr231^ (1:1000; AT180, Thermo Scientific, USA). The secondary antibodies used against above were anti-Mouse Cy3 (1:200), anti-Mouse Alexa 488 (1:200), anti-Rabbit Alexa 488 and anti-Rabbit Cy3 (1:200) procured from Molecular probes, USA.

### Protein extraction and western blotting

For protein extraction, homogenization of decapitated heads of adult flies was performed in lysis buffer, prepared by 0.05 M Tris-Cl (ph 8.0), 0.1% SDS, 0.15 M Nacl, 0.02% Na-Azide, 1% Triton X-100, 0.5% Na-deoxycholate, phosphatase inhibitor cocktail (Thermo Scientific, USA) and protease inhibitor cocktail (Thermo Scientific, USA). The total protein amount was estimated by Pierce^R^ BCA Protein Assay Kit (Thermo Scientific, USA) and equal amount of it was loaded and separated by SDS-PAGE, followed by transfer onto a PVDF membrane (Millipore, USA) for overnight at 4°C. Consecutive day, membrane was washed, blocked using the Pierce^R^ Protein-Free (TBS) blocking buffer and probed with primary antibody at 4°C for overnight.

The primary antibodies utilized were anti-human tau (5A6; 1:1000, DSHB, USA), anti-human PHF-tau^Thr231^ (AT180; 1:1000, Thermo Scientific, USA), PHF-tau^Thr181^ (AT270, 1:1000, Thermo Scientific, USA), anti-GFP (4B10; 1:1000, Cell Signalling technology, USA) and anti-β-tubulin (E7; 1:1000, DSHB, USA). The primary antibodies were detected by secondary antibody conjugated to horseradish peroxidase (HRP) (1:2000, GeNei™, India). The blots were developed in SuperSignal^R^ West Pico Chemiluminescent Substrate (Thermofisher Scientific, USA) and imaged using Fujifilm LAS 4000. The blots were at least triplicated in every experimental setup and densitometric analysis of each was performed through Image J software.

### Imaging and statistical analysis

Capturing of images was done using Leica TCS-SP5 II and/or SP8 Falcon confocal microscope. The images were processed and assembled using the Leica application suite advanced fluorescence software and Adobe Photoshop CS5 software respectively. MS Excel 2013 and/or GraphPad Prism 8.3.1 was utilised for graphical representation of the data. The errors bars/ ± sign denotes standard deviation in the data set. For statistical analysis, paired *t* test and One-Way ANOVA was applied to the data, and p-values <0.05 were considered as significant, with p-value ≤ 0.05 as *, p-value ≤ 0.01 as ** and p-value ≤ 0.001 as ***.

## Results

### Human tau doesn’t show trans-cellular propagation at the *Drosophila* larval stage despite of robust tau expression

*Drosophila* models of tauopathies have shown to confer neurodegeneration using various specific drivers of the *UAS-Gal4* system [17]. Among which, Glass multiple reporter (GMR), *GMR-Gal4*, is one of the widely accepted and used to study the neurodegenerative disease aetiology in eyes [16, 18, 19]. We have utilized *GMR-Gal4* to drive the expression of wild type form of human tau (h-tau^WT^) and green fluorescence protein (GFP) transgene and examined the nature of tau distribution in terms of probable trans-cellular migration.

As the schematic diagram in Fig 1. F, depicts, in the larval stage, *GMR-Gal4* expresses in the differentiating photoreceptor cell (PRC) of the morphogenetic furrow in the eye-antennal discs and their axons projecting into the optic lobes of the brain via the optic stalks [18, 20].

**Figure 1.**
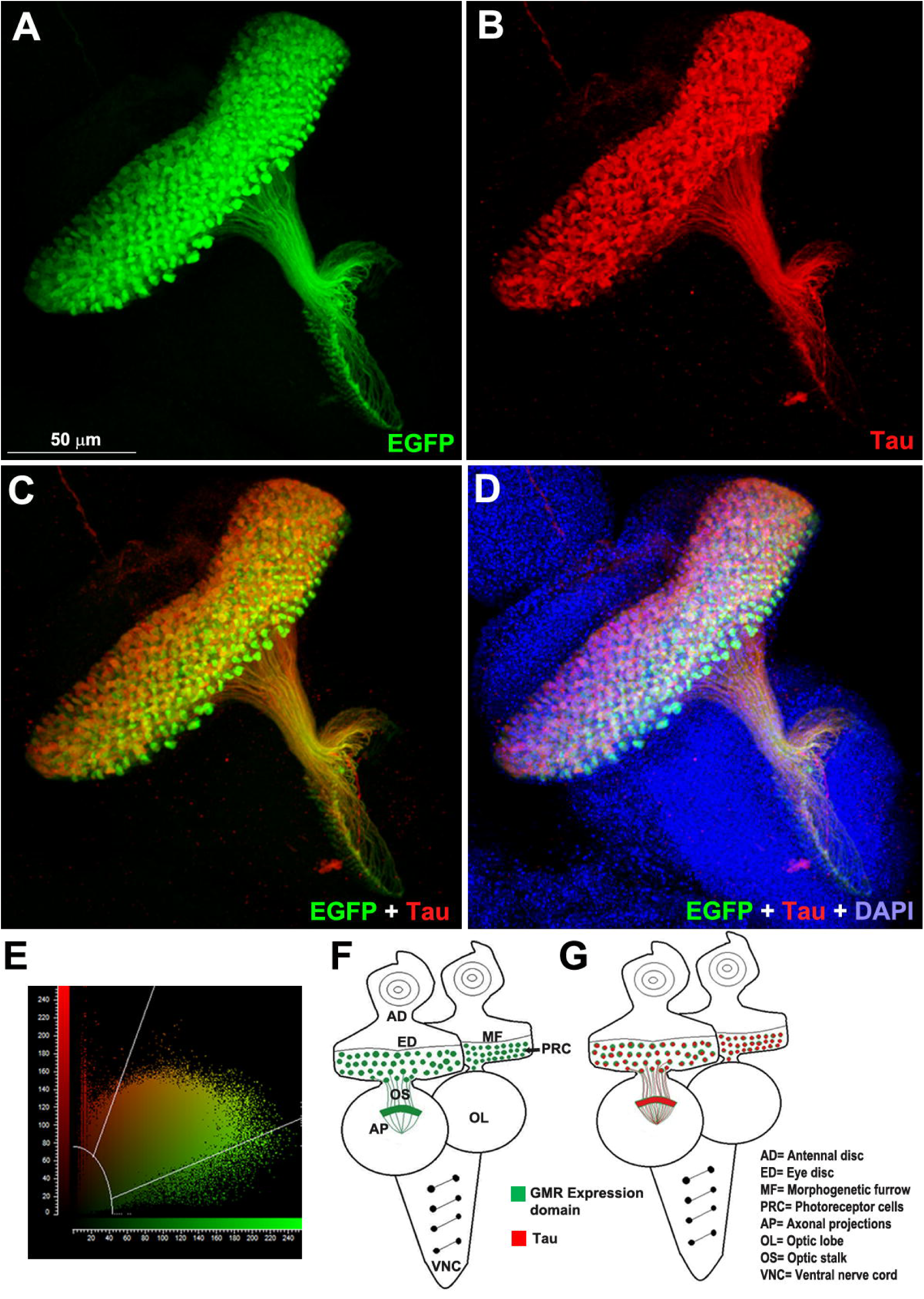
Wild type form of human tau does not show migratory property in *Drosophila* larval tissues regardless of its prominent expression. (A) Expression pattern of *GMR-Gal4* as marked by EGFP (green) in the third instar *Drosophila* larval eye disc and brain. (B) Expression pattern of *GMR-Gal4* driven human tau^WT^ transgene. (C, D) Near complete co-localization of GFP (green) and tau (red) suggests immobility of the tau protein beyond the Gal4 domains. (E) A generated scatter plot of the image C confirms an average of 90.83% co-localization between green (EGFP) and red (tau) signals. (F) Schematic representation of the *GMR-Gal4* expressing domains in the larval eye disc and brain. (G) Schematic showing the relative distribution pattern of GFP and tau in the corresponding *GMR-Gal4* > *GFP* + *tau*^*WT*^ larval tissues. DAPI (blue) used a counter stain in D. Scale bar in A-D= 50 µm.

Further in the direction of our hypothesis, larval eye discs and brain tissue of the flies co-expressing *GMR-Gal4* driven transgenes GFP and h-tau^WT^ (*GMR-Gal4* > *GFP* + *tau*^*WT*^), were analysed for the endogenous GFP expression and relative tau distribution (Fig 1. A-D). The reporter gene, GFP was utilized to mark the *GMR* expression domain. In line with the previous reports [20], endogenous GFP was attained in the destined region, below the morphogenetic furrow and axons connecting it to the brain (Fig. 1 A). This confirms that tau expression has no impact on the *GMR* expression in terms of its area.

While the expression of GFP as predicted, was limited to the *GMR* expression zone, the expression of tau also co-localized within the GFP domain, as represented by the image and corresponding schematic in Fig. 1 C and G respectively. The co-localization coefficient between both the ROI (region of interest), eGFP: tau, was estimated using Leica application suite advanced fluorescence software (LAS AF Lite) [21] and was calculated to be 90.83%, with the remaining percentage corresponding to the non-specific outlier data, as precisely depicted in the scatter plot in Fig. 1 E. The above observations suggest that, although the larval tissues exhibit a notable expression of tau throughout the *GMR* domain (Fig. 1 B), however, their propagation or movement is not ensued in the larval tissues.

### Human tau propagates trans-cellularly in adult *Drosophila* by the factor of age

Following the larval tissues, which were found to be naive with respect to tau propagation, the tau migration at adult stages was scrutinized. The formation of tau aggregates in forms of paired helical filaments (PHFs) and NFTs have already been reported in the adult flies [19, 22]. To investigate if these entities possess any trans-cellular migratory nature in *Drosophila*, we analysed the relative expression domains of the *GMR* driven GFP and h-tau^WT^ transgenes in the aging adult heads.

As illustrated in the schematic in Fig. 2 H, the expression pattern of *GMR* has been found to be confined to the retina, lamina and axonal terminals of the photoreceptors reaching their target zones in parts of the medulla [20, 23]. Here, it also important to note that *GMR-Gal4* has already been utilized to demonstrate the transcellular spreading of huntingtin aggregates in the adult fly brain [24].

**Figure 2.**
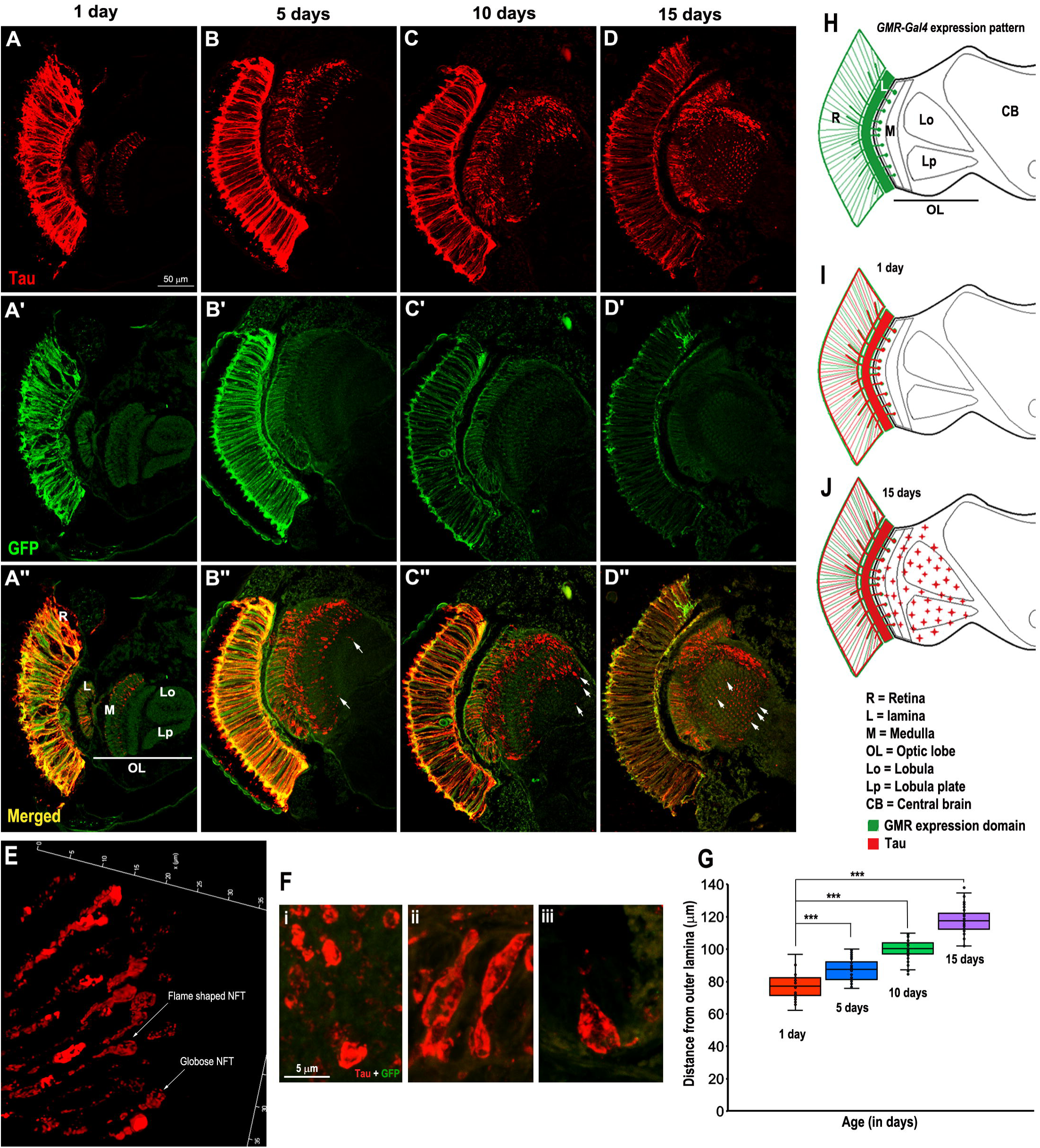
Age dependent trans-cellular propagation of h-tau^WT^ across *GMR-Gal4* > *GFP* + *tau*^*WT*^ adult *Drosophila* head. Human tau aggregates (red in A-D) migrate beyond the expression domains of *GMR-Gal4* marked by GFP (green in A’-D’) in aging adult flies. (A□’-D□’) The merged images depict tau expression within the Gal4 boundaries at day 1 (A”), while a few aggregates migrate beyond the zones of the *GMR-Gal4* at day 5 (arrows in B”), and with an increase in the distance on day 10 (arrows in C”). The migration continues and the majority of the tau aggregates cover the optic lobe by 15^th^ day of aging (arrows in D”). 3D view of tau aggregates captured at high resolution lighting mode (E) and in conventional mode (F) revealed formation of highly prominent globose (marked in E; F-i) and flame shaped (marked in E; F-ii, iii) neurofibrillary tangles. The distance of tau aggregates from the outer boundary of the lamina is quantified at various aging points and represented as a box plot (G). Schematic representations of the adult head section of *Drosophila* illustrating the expression domain of *GMR-Gal4* (H), relative expression of tau and *GMR* in 1 day and 15 days old experimental tissues (I and J respectively). ^***^*P*<0.001 using student t-test and One-Way ANOVA. Scale bar A-D” = 50 µm, F (i-iii) = 5 µm.

The adult flies co-expressing GFP and h-tau^WT^ transgenes (*GMR-Gal4* > *GFP* + *tau*^*WT*^) were utilized with the motive to mark the *GMR* expression domain by the reporter protein-GFP, and drive the disease in the same region. Therefore, if tau will possess any tendency to propagate, it could be conveniently analysed in the adult head sections. Accordingly, the propagated regions will be those domains of the adult head, where the expression of tau would be evident while that of reporter gene would be absent. The adult flies of the *GMR-Gal4* > *GFP* + *tau*^*WT*^ genotype were aged for 1, 5, 10 and 15 days to investigate the GFP expression and relative distribution pattern of tau aggregates in the head sections. To maintain the uniformity, the tissue sections comprising the middle portion of the adult heads were chosen for analysis. The head sections were comprehensively examined and analysed by confocal microscope, Leica TCS SP5 II at 20X with 1.8 digital zoom (Fig. 2 A-D”) and 40x with 2.0 digital zoom (Fig. S2).

In agreement to the earlier reports [20, 23], the expression of the GFP was restricted to the predicted *GMR* region, with robust expression in the retina, lamina and in the axonal terminals of the outer medulla in 1 day old flies (Fig. 2 A’; arrowhead mark the axonal terminals in Fig. S2 A’). The tau expression was also marked in the similar regions on 1day post ecolsion, along with the tau aggregates at the axonal terminals in the outer medulla of the optic lobes (Fig. 2 A; Fig. S2 A). Intriguingly, during the course of aging these tau aggregates were seen to migrate from the axonal terminals of the outer medulla to the other regions of the brain where the *GMR-Gal4* does not express and the subsequent GFP expression was absent (Fig. 2 A”-D”; Fig. S2 A”-D”). The migration of these aggregates was measured in the term of distance travelled by them from the outer boundary of the lamina. As depicted in Fig. S1 A-D, for every head section, the distance was measured across the mid-point of the outer boundary of the lamina to the farthest propagated aggregate from that particular point to maintain the uniformity while data collection and analysis. Hence, some of the aggregates those had migrated beyond the calculated distance, but were not in line with the mid-point of the outer boundary of lamina (Fig. S1 arrowhead in B) were excluded from distance quantification, and therefore, the actual propagated distance in some tissues was even more than the estimated value.

In 1 day old adult flies, the formation of aggregates was witnessed till the outer medulla of the brain (*GMR* expression domain), for about 76.95±7.26 µm from the lamina, with almost a complete co-localization between GFP and tau expression (Fig. 2 A, A”; Fig. S2 A, A”; correlate with schematic in Fig. 2 I). A majority of the tau aggregates found at day 1 were restricted to the expression domain of *GMR-Gal4* (Fig. S2 A”, arrows), except rare events where a few of such tau entities were also noted outside the boundaries of the Gal4. Since majority of the tau aggregates were confined within the limits of *GMR* expression domain on day 1, thereby, this was considered to be the baseline for the comparative analysis of tau propagation during subsequent aging.

Intriguingly, in 5 days old flies, a few of the tau aggregates were found to migrate from the axonal terminals in the *GMR* expression domain and spread towards the other regions of the optic lobe in the brain, travelling upto an average distance of 87.42±6.76 µm from the lamina (arrows in Fig. 2 B”; arrow heads in Fig. S2 B”; also see Fig. S1 B). Perhaps, the migration potency is ensued between the day 1 and 5 in adult flies. Further, the distance travelled by tau aggregates and their cellular abundance tends to increase with age, as in 10 days old flies, these entities were observed to move up to 99.9±6.15 µm distance from the lamina. (arrows, Fig. 2 C”, S1 C; arrow heads, S2 C”). Subsequently, in 15 days old flies, the degree of trans-cellular propagation was found to be maximum, where a majority of the aggregates were visualised outside the *GMR* expression domain with up to an average distance of 117.79±8.27 µm from the lamina (arrows, Fig. 2. D”; S1 D; arrow heads, S2 D”, correlate with schematic in J). The optic lobe of such brain was found to be almost covered by the propagated tau aggregates (arrows in Fig. 2. D”; Fig. S1 D; arrowheads in Fig. S2 D”). Notably, the tau aggregates migrate an average of 40.84 µm distance in adult brain from 1 day to 15 days of aging. Interestingly, the 3D projection images captured by analysing the tissues from 15 days old flies at high resolution scale of lighting mode using Leica TCS SP8-Falcon confocal microscope revealed robust presence of flame shape and globose NFTs (arrows in Fig 2. E). Additionally, Fig. 2F depicts the globose (Fig. 2 F i) and flame shaped (Fig. 2 F ii, iii) morphology of the migrating tau aggregates. Comprehensive representation of the data set across aging clearly establishes a positive correlation between the migration potency of the tau aggregates and aging (Fig 2 G). It is also worth mentioning here that in comparison to the tau signals, the GFP signals was progressively lost in the axonal terminals with age, which seems to be due to the subsequent degeneration of the axonal neurons/terminals (Fig. S2 A’-D’). Therefore, above experiments distinctly demonstrate age-dependent trans-cellular propagation of h-tau aggregates in adult *Drosophila* head.

### An additional transgene doesn’t induce the propensity of tau to propagate beyond trans-cellular boundaries

Further, we investigated if introduction of an additional transgene in the *GMR-Gal4* > *tau*^*WT*^ flies (i.e. *UAS-GFP* transgene in the previous experiments) had persuaded any trans-cellular propagation prophecy in the tau aggregates. To examine this, we traced the migratory behaviour of the tau aggregates in 1 day and 15 days old flies which solely expressed *GMR-Gal4* driven h-tau^WT^ transgene. The distance travelled by the tau aggregates in *GMR-Gal4* > *tau*^*WT*^ flies was estimated in similar manner as performed with the *GMR-Gal4* > *GFP* + *tau*^*WT*^ flies (Fig. S1).

The head sections of the age matched *GMR-Gal4* > *tau*^*WT*^ flies revealed comparable migratory patterns of the tau entities between both the genotypes (*GMR-Gal4* > *tau*^*WT*^ flies with or without *UAS-GFP*). As noted in the *GMR-Gal4* > *GFP* + *tau*^*WT*^ flies, the existence of tau aggregates in the head section of 1 day old *GMR-Gal4* > *tau*^*WT*^ flies was evident in the retina, lamina and outer medulla, with an average distance upto 102.02±9.22 µm from the lamina (compare Fig. S3 A with Fig. 2 A, A”). Interestingly, in 15 days old *GMR-Gal4* > *tau*^*WT*^ adults, the tau aggregates had migrated to spread onto cover the optic lobes of the brain (compare Fig. S3 C with Fig. 2 D, D”), with an average travelling distance upto 133.31±10.51 µm (compare Fig. S3 A, C). Quantitative analysis suggests that the tau aggregates travel an average distance of 31.3 µm in *GMR-Gal4* > *tau*^*WT*^ adult heads during the 15 days of aging (Fig. S3 E). This outcome further wires the concept that, phenomenon of tau migration across trans-cellular borders in fly model is not ensued by the presence of any additional reporter gene (GFP transgene in the present study) in its genome, rather is an autonomous and intrinsic property possessed by the tau aggregates.

### Hyper-phosphorylated tau increases quantitatively and migrates trans-cellularly over age

The pathogenic identity of tau is attained due to various factors, among which its hyper-phosphorylation one such mechanism [3]. Physiological tau contains about 2-3 moles of phosphate/mole of tau protein [25, 26]. The homeostasis of phospho tau is maintained by kinases and phosphatases [27]. However, in its pathogenic form, tau phosphorylation is enhanced, subsequently deteriorating its biological activity, namely stabilization and assembly of microtubules [27, 28]. Consequently, tau detaches itself from the microtubules and tends to aggregate as NFTs [29].

To determine if there is any correlation of hyper-phosphorylation of tau to its propagation, we considered examining it in our model with respect to aging. Among the approximately 40 studied tau sites that are abnormally phosphorylated by a plethora of kinases in AD, site Thr 231 (recognised by AT180 antibody) was selected [27, 30]. The abundance of phosphorylated tau at this particular epitope was examined in *GMR-Gal4* > *tau*^*WT*^ flies, aged for 1, 5, 10 and 15 days. Although similar propagation properties of tau were observed in both the genotypes i.e. *GMR-Gal4* > *GFP* + *tau*^*WT*^ and *GMR-Gal4* > *tau*^*WT*^, yet we chose *GMR-Gal4* > *tau*^*WT*^ flies for further experiments.

The state and nature of the phospho-epitope analysed by staining in the 1 day old adult head sections revealed robust presence of phosphorylated tau in the retina, lamina and a part of the optic lobe in the form of aggregates (Fig. 3 A). Interestingly, the aggregates manifested substantial levels of phosphorylation, even in the 1 day old flies. Though, almost all of those entities (Fig. 3 A) were limited to the outer medulla (expression domain of *GMR-Gal4*), approximately at 91.24±16.70 µm distance from the lamina, except a few cases in which the phospho-tau aggregates appeared to escape the Gal4 expression domain. However, by the age of 15 days, the presence of phosphorylated tau aggregates was apparent beyond the trans-cellular boundaries of Gal4 domain. Following a similar pattern as noted earlier (Fig. 2 D”), the tau aggregates showing prominent phosphorylation were seen to move towards the central brain, travelling upto 127.16±9.40 µm from the lamina (Fig 3. D). Quantitative analysis reveal that phospho-tau aggregates travel an average distance of 35.92 µm during the 15 days of aging (Fig. 3 E).

**Figure 3.**
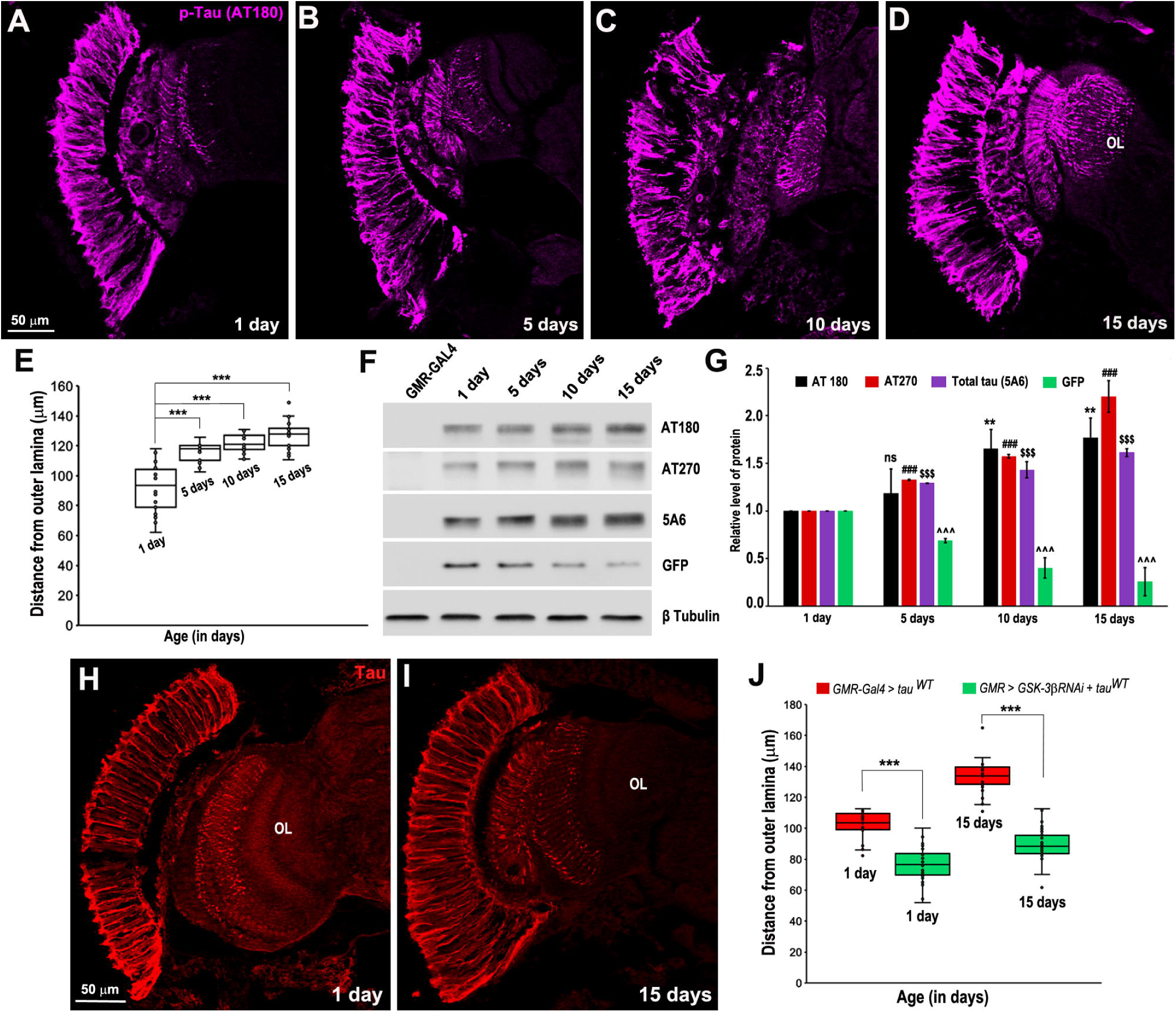
Age dependent trans-cellular migration of tau aggregates shows a positive correlation with phosphorylation, which could be restricted by down-regulation of *Gsk-3β*. (A) In 1 day old adults (*GMR-Gal4* > *tau*^*WT*^), majority of the tau aggregates those are restricted within the *GMR-Gal4* expression domains are phosphorylated. (B-D) By the 15^th^ day of aging, the hyperphosphorylated tau aggregates cross the Gal4 boundary and cover the additional areas the optic lobe. (E) Graphical representation of the distance of phospho tau aggregates from the outer boundary of the lamina across aging. (F) Western blots of the lysates isolated from aging adult heads (*GMR-Gal4* > *tau*^*WT*^) showing progressive increase in the level of tau phosphorylation at pT231 site, pT181 site and tau titre, as detected by AT 180, AT 270 and 5A6 antibodies, respectively. Comparative reduction in the levels of GFP over aging in *GMR-Gal4* > *GFP* + *tau*^*WT*^ adult flies. (G) Quantification of phospho tau, total tau and GFP normalised to β tubulin. (H-J) Targeted downregulation of *Gsk-3β* restricts the migration propensity of tau aggregates in aging adult heads. (H) In 1 day old adults, the tau aggregates can be visualised within the domains of *GMR-Gal4*. (I) The tau aggregates which generally migrate to cover the optic lobe in 15 days old adult *GMR-Gal4* > *tau*^*WT*^ flies exhibited restricted migration upon downregulation of *Gsk-3β*. (J) Comparative graphical representation of the distance of tau aggregates from the outer boundary of the lamina with the course of aging in *GMR-Gal4* > *tau*^*WT*^ and *GMR-Gal4* >*Gsk-3β* + *tau*^*WT*^ adult flies. \*\**P*<0.01, ^***^*P*<0.001 using student t-test and One-Way ANOVA, wherever applicable. OL = optic lobe; Scale bar A-D, H-I= 50 µm.

Further, the amount of phospho-tau was examined by western blot analysis by AT180 antibody, and also verified by another phosphor-tau specific AT270 antibody [31] in the protein isolated from 1, 5, 10 and 15 days old adult fly heads (Fig. 3 F). The relative level of phosphorylated tau as examined by both the antibodies was found to increase from 1 day to 15 days old flies (*GMR-Gal4* > *tau*^*WT*^), thereby validating the fact that the amount of phospho-tau also increases with age (Fig. 3 F) as does the migration of tau aggregates (Fig. 2 A”-D”). Here it is important to note that the relative levels of total tau were also found to increase by the factor of age in *GMR-Gal4* > *tau*^*WT*^ flies (Fig. 3 F). Such increase in the level of total tau can be articulated to the stalled removal of tau aggregates due to impaired clearance machinery in the disease condition [32, 33]. To further validate the above, we checked the levels of GFP in the age matched *GMR-Gal4* > *GFP* + *tau*^*WT*^ adults, and interestingly, the levels of the protein was noted to decline over age (Fig. 3 F). This suggests that age dependent increasing level of phosphorylated and total tau is not a generalized phenomenon. Statistical analysis of the densitometric data generated from the 3 independent western blot experiments, provides a comparative depiction of the age dependent increasing level of phosphorylated and total tau when normalized to β tubulin (Fig. 3 G)

From the above observations, it can be articulated that though abundance of the phosphorylated tau increases over aging, however, it is yet difficult to establish the exact involvement of phosphorylation in trans-cellular tau migration.

### Down-regulation of Gsk-3β kinase restricts trans-cellular migration of tau aggregates

Among the variety of reported kinases that phosphorylate tau, Glycogen synthase kinase-3β (Gsk-3β) phosphorylates approximately 36 residues in the tau protein, which also include the above examined Thr 231 epitope [27, 34]. Previous reports have shown that this priming kinase when co-expressed with tau transgene exaggerates the neurodegeneration in fly models [35]. In contrast, its down-regulation by inhibitors has also shown to rescue the tau mediated disease symptoms [36]. This makes GSK-3β a promising candidate to investigate if its down-regulation causes any impact on age dependent tau migration in the adult fly heads.

Accordingly, tau migration was evaluated in the tissue sections of *GMR-Gal4* > *Gsk-3β RNAi* + *tau*^*WT*^ flies aged for 1 day and 15 days. In agreement to the earlier reports [36], we too noted that downregulation of Gsk-3β causes a significant rescue in terms of neurodegeneration in the internal architecture of adult eyes (Fig. 3 H-I). Staining experiments in the head sections of 1 day old *GMR-Gal4* > *Gsk-3β RNAi* + *tau*^*WT*^ flies revealed the presence of tau aggregates in the retina, lamina and in parts of optic lobe of the brain, with average distance of 76.80±11.13 µm from the outer boundary of lamina (Fig. 3 H). Here it is important to note that the tau staining pattern (retina, lamina and aggregates in the outer medulla) in 1 day old *GMR-Gal4* > *Gsk 3β RNAi* + *tau*^*WT*^ was almost comparable to the pattern observed in the age matched *GMR-Gal4* > *tau*^*WT*^ flies (compare Fig S3 A with Fig. 3 H). Intriguingly, however, propagation of tau aggregates which was common in 15 days old *GMR-Gal4* > *GFP* + *tau*^*WT*^ (Fig. 2 D, D□□) or *GMR-Gal4* > *tau*^*WT*^ flies (Fig. S3 C), was essentially restricted by the down-regulation of *Gsk-3β*, with the aggregates showing an average distance of 89.95±11.26 µm from the lamina (Fig. 3 I). The box plot shows the trends in the age dependent migration of tau aggregates in *GMR-Gal4* > *tau*^*WT*^ flies with and without *Gsk-3β RNAi* transgene expression (Fig. 3 J). The tau aggregates at day 15, that had eventually migrated and covered the optic lobe in *GMR-Gal4* > *tau*^*WT*^ flies, reaching upto an average of 31.30 µm away from the borders of the GMR domain, were evidently observed to be constrained upon downregulating *Gsk-3β*, with travelling upto only an average of 13.15 µm distance (Fig. 3 J). Therefore, a significant rescue in terms of tau migration is evident in flies of *GMR-Gal4* > *Gsk-3β RNAi* + *tau*^*WT*^ genotype as compared to *GMR* > *tau*^*WT*^ flies.

It can be scrutinized from the above that the migration potential of tau aggregates is compromised with the down-regulation of *Gsk-3β*, which actually restores the phosphorylation status of tau [27]. This also point to the possibility of enhanced tau phosphorylation over aging, being involved in driving the migration of tau entities. Hence, hyper-phosphorylation of tau perhaps instigates the trans-cellular tau migration.

### The intrinsic property of human tau to propagate is retained in another tau^V337^ isoform

We next investigated if the migratory nature of tau aggregates is also persisting in additional isoform of tau protein. The *Drosophila* transgenic flies, *UAS-tau*^*V337*^ which expresses the mutated version of tau (Val^337^→Met), reported to be associated with early onset and familial dementia [37, 15], and *GMR-Gal4* driven expression caused degeneration of the eyes, as reported previously [15, 19].

The localization pattern of tau aggregates was evaluated in the adult head sections of *GMR-Gal4* > *tau*^*V337*^ flies aged for 1 day and 15 days. At day 1, robust expression of tau was retained in the retina and lamina, and tau aggregates were apparent in the outer medulla of the optic lobe of the brain (Fig. 4 A). Interestingly, the migration characteristics of the tau aggregates in *GMR-Gal4* > *tau*^*V337*^ flies was found to be similar as noted with *h-tau*^*WT*^ allele. Compared to the tau aggregates those were mostly confined to the *GMR* expression zone in 1 day old *GMR-Gal4* > t*au*^*V337*^ flies, the aggregates migrated beyond the Gal4 expression domain and travelled to cover additional areas of the optic lobe in 15 days old flies (arrowheads in Fig. 4 B). The box plot in Fig. 4 C represents an increasing trend in distance of tau aggregates from outer boundary of the lamina when estimated across aging. Henceforth, the ability of tau to spread trans-cellularly is not dependent on its isoform.

**Figure 4.**
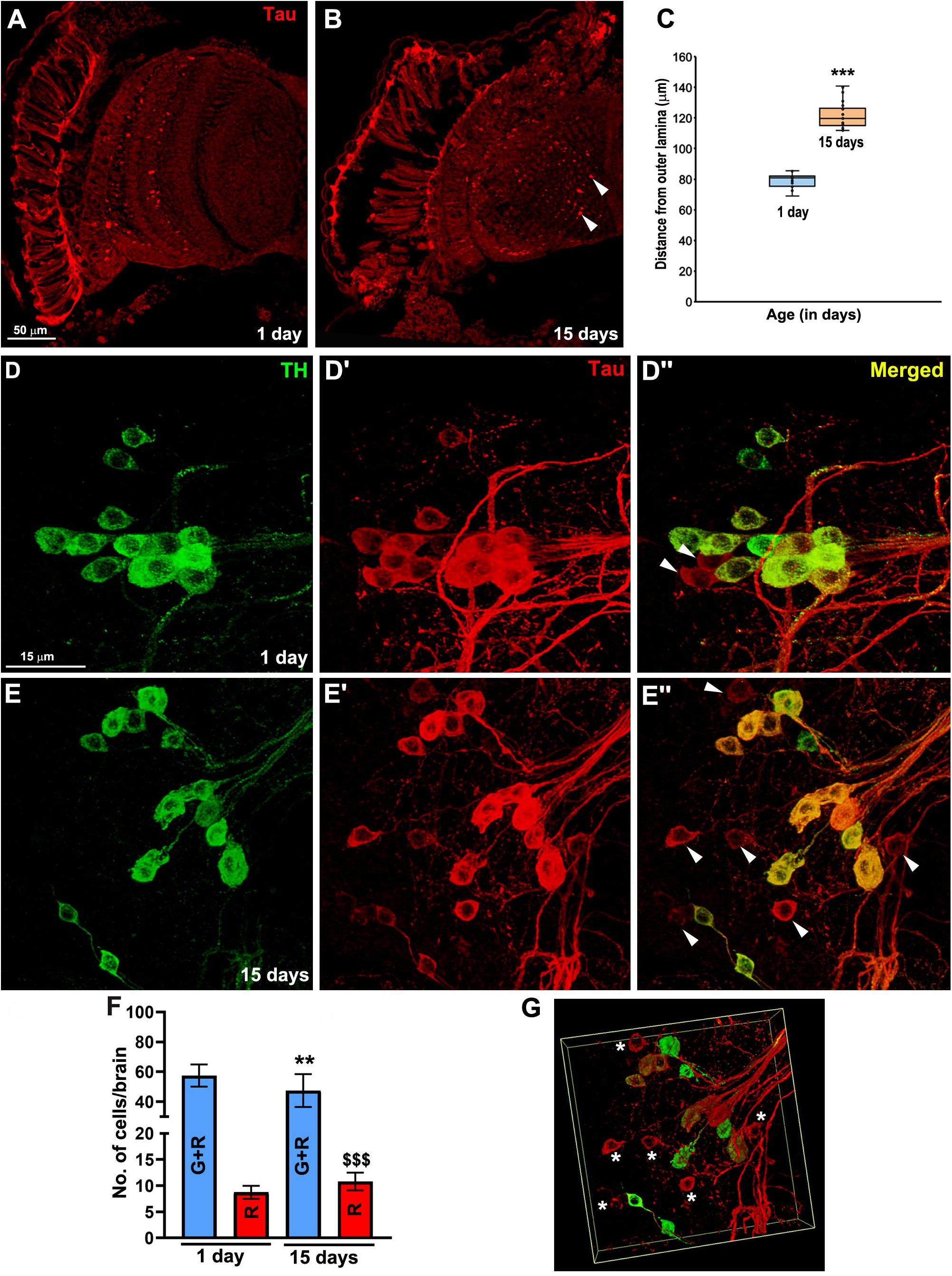
Intrinsic property of tau aggregates to migrate across the trans-cellular borders is retained in additional h-tau isoform, and Gal-4 driver. (A) In 1-day old head sections of *GMR-Gal4* > *tau*^*V337*^ adult flies, tau expression (red) was confined to the domains of *GMR-Gal4* expression. (B) In 15 days old adult head sections, migration of tau aggregates was apparent beyond the Gal4 domain and reached upto the additional areas of the optic lobe of the brain (arrow heads). (C) Graphical representation of the distance of tau aggregates from the outer boundary of the lamina as a function of aging is represented as a box plot. (D, E) In *TH-Gal4* > *tau*^*WT*^ adults, the expression domain of *TH-Gal4* is marked by anti-TH staining (green). (D’, E’) Expression pattern of *TH-Gal4* driven human tau^WT^ transgene (red). (D”, E”) Merged images of TH and Tau staining. (D-D”) In 1 day old adults, compared to the total number of cells with both the TH and tau expression, only a few of the tau positive identities were devoid of any TH expression (arrow heads in D”). (E-E”) 15 days of aging caused an increase in the number of tau positive cells and subsequent decline in the number of cells with co-localized TH and tau expression (arrowheads in E”). (F) Comparative representation of the relative numbers of cells per brain with TH and tau (G+R) and only tau expression (R). (G) 3D projection image of E” recaptured in high resolution lightning mode to validate the increased abundance of the cells with tau aggregates (marked by asterisk). ^**^*P*<0.01, \*\*\**P*<0.001 using student t-test. Scale bar A-B= 50 µm, D-E” = 15 µm.

### The trans-cellular migration potency of tau is preserved with another Gal4 line: *TH-Gal4*

In addition to a different tau isoform, we also validated the migratory characteristics of the tau aggregates by an additional Gal4 driver line, the *TH-Gal4* (tyrosine hydroxylase-Gal4) which expresses specifically in the dopaminergic neurons of the brain [38]. The dopaminergic system of the adult brain comprises of the cluster of neurons studied and validated in several previous studies [22, 38, 39]. We examined the comparative expression of TH and Tau in the clusters of dopaminergic neurons of 1 and 15 days old adult brains of *TH-Gal4* > *tau*^*WT*^ flies.

In line with the previous reports [22, 39], the expression of TH (green) was observed in the clusters of the dopaminergic neurons (Fig. 4 D), and the tau (red) followed the similar expression pattern in 1 day old *TH-Gal4* > *tau*^*WT*^ brain (Fig. 4 D’). It is increasingly known that the expression of human tau in dopaminergic neurons form NFT like tau aggregates [22], and causes progressive degeneration of the neurons. In line with the previous reports, we also observed progressive degeneration of dopaminergic neurons in *TH-Gal4* > *tau*^*WT*^ flies approximately post 2 weeks of eclosion. In 1 day old adults, the expression of tau mostly co-localized with that of TH, with some tau positive cells devoid of TH expression (arrowheads in Fig. 4 D”). With the increasing age of the fly, (till day 15) the tau aggregates appear to migrate in those cells which were devoid of any *TH-Gal4* expression, thereby increasing the number of the only tau positive cells (arrowhead in Fig. 4 E”). Hence, it is interesting to note that the numbers of tau positive cells without any TH expression exhibited a positive correlation with increasing age (Fig. 4 F). For further validation, 3D projection image of the 15 days old brain area was captured and analysed from various angles, one of which is represented in Fig. 4 G, and propagated tau aggregates are marked by asterisks. Therefore, above observation further demonstrates the *Gal4-*driver independent inherent trans-cellular propagation property of h-tau aggregates in *Drosophila* adult brain.

## Discussion

The potency of the misfolded tau to migrate and spread across the brain is addressed to be one of the major causes of the pathological manifestations and aggravation of the tauopathies in human. It is quite evident from the previous reports that tau aggregates do migrate across trans-cellular borders [6, 40]. These previous studies have utilized various model systems to elucidate this phenomenon, which are either *in-vitro* or *in-vivo* in nature, with former not being able to replicate the actual biology of the human brain [41]. While the *in-vivo* systems, either involve infusion or injection of the exogenous tau aggregates [13], or those utilizing endogenous tau expression to study its migration consume long incubation periods to develop the propagation pathology [42, 43]. Henceforth, the present hour demands the creation of such *in-vivo* model systems that allow rapid and convenient study of this aspect of disease aetiology as in-depths of the mechanism are still a prime area of neurobiology research.

The *Drosophila* tauopathy model(s) recapitulates the pathological determinants of the disease in terms of neurodegeneration and aggregate formation. Previous reports have claimed the formation of tau aggregates in forms of PHFs and NFTs in the adult flies [19, 22]. We tested if these pathogenic tau aggregates possess the similar tendency of trans-cellular migration in *Drosophila* as they are reported in the higher organisms and in *in-vitro* studies as stated above. Fly disease models offer innumerable advantages over other model systems such as rapid disease onset, easy phenotypic scoring in bulk, ease to study the status of nervous system, and to elucidate the effect of molecules and genetic modifiers on disease pathogenesis and phenotypic manifestation [12].

Previous studies have already demonstrated that nervous system specific expression of human tau in *Drosophila* larvae causes disruption of axonal microtubule cytoskeleton and development of locomotor dysfunction [44, 45]. Therefore, the initial evaluation of the probable migratory potency of tau aggregates at larval stage was scrutinized, however, no transcellular propagation was noted, despite of prominent expression of tau in the target tissues. Hence, the larval stages could be designated as early phase of tauopathy manifestation. Subsequent analysis in the adult flies revealed trans-cellular propagation of tau aggregates in aging brains. Here, it is worth noting that since *GMR-Gal4* expression is well documented and has already been utilized in several studies [20, 23, 24], and therefore, propagation potency of tau aggregates to migrate and spread across the brain in our model is not due to the leakiness of the Gal4 promoter. Also, the tau propagation observed in the present study is independent of tau isoforms and the utilized Gal4 drivers. The absence of spreading of tau aggregates at the larval stage, while its subsequent propagation in the adults is probably due to the lack of disease progression at the former stage. Interestingly, the analysis of tau migration potency being age dependent in our data is in line with the previous reports with other model systems [14]. It appears that tau requires a course of maturation and increase in its titre to achieve the trans-cellular migratory potential, which happens over time with aging.

Though the role of phosphorylation in tau aetiology is well documented [27, 46], however, at present it is difficult to establish a direct correlation between age dependent increase in the tau propagation and enhanced phosphorylation in *Drosophila*. Though, the restricted migration of tau aggregates due to down-regulation of *Gsk-3β* did suggests an involvement of phosphorylation in acquiring the migratory property. It appears that tau hyperphosphorylation over aging progressively increases the load of toxic tau species in the target tissues, which then spread to the newer cells/regions. Hence, regulation of tau phosphorylation can be exploited as an effective strategy to restrict subsequent propagation of tau entities in unaffected regions of the brain. Moreover, our model provides a unique opportunity to identify and characterize novel regulators of tau propagation.

Taken together, for the first time we demonstrate age dependent trans-cellular migration of h-tau in *Drosophila*. Moreover, our findings have developed a *Drosophila* based endogenous h-tau expressing novel *in-vivo* model with immense application potential that successfully recapitulates the trans-cellular migration potency of tau aggregates. Our system provides an easy, rapid and quantifiable evaluation of the cellular and molecular insights which facilitate this astounding biological phenomenon. Hence, the present study can be immensely helpful in designing novel approaches to restrict the transcellular spreading of tau aggregates, which has been considered as a key factor of the disease aetiology.

## Acknowledgments

We are thankful to Prof. Mel Feany (Harvard Medical School, USA), Bloomington Stock Center and Developmental Studies Hybridoma Bank (DSHB), USA, for providing some fly stocks and antibodies used in this study. This work is supported by a research grants [Ref. no. BT/010/IYBA/2017/02] from the Department of Biotechnology (DBT), Government of India, New Delhi, India, to SS. Ms. Aqsa is supported by Junior Research Fellowships (JRF), from Council of Scientific and Industrial Research (CSIR), New Delhi, India. We are grateful to Ms. Nabanita Sarkar for the technical support.

## Supplementary figure legends

**Figure S1.** Standard parameter applied for quantification of distance of tau aggregates from outer boundary of the lamina in aging adult heads for all experiments. The distance of tau entities was measured from the middle point of outer boundary of the lamina to the farthest propagated aggregate in the direction of that reference point, subsequently depicted via a white dotted line in A-D. Mentioned numbers are the distance of tau aggregates for that particular tissue section. Arrowhead in B shows a migrated tau aggerate which is not in line with the applied quantification parameter.

**Figure S2.** Analysis of head sections of *GMR-Gal4* > *GFP* + *tau*^*WT*^ adults at higher magnification (40x + 2 digital zoom) reveal the in-depth pattern of the age dependent migration properties of tau aggregates. (A-D) *GMR-Gal4* driven tau distribution (red) across aging adult heads. (A’-D’) The expression domain of *GMR-Gal4* marked by GFP is retained in the retina, lamina and axonal terminals (arrowheads) in the outer medulla. The GFP expression is progressively lost with aging. (A”) In 1 day old adults, tau expression can be noticeably observed within the expression zones of *GMR-Gal4* (arrows). (B”) In 5 days old adults, some tau aggregates migrate across the trans-cellular borders of *GMR-Gal4* to localize in the additional areas of the brain (arrowheads). (C”) By 10^th^ of aging, tau aggregates travel an additional distance beyond the transcellular boundary of Gal4 domain (arrowheads). (D”) Maximum propagation is observed in 15 days old adult heads where a majority of the tau aggregates cover additional regions of the optic lobe of the brain (arrowheads). Scale bar A-D”= 50 µm.

**Figure S3.** The intrinsic migratory property of tau aggregates is not a consequence of an additional *UAS-GFP* transgene in the genomic background of *GMR-Gal4* > *tau*^*WT*^ flies. (A-B) Robust expression of tau is limited within the *GMR-Gal4* expression boundaries in 1 day old adult head. (C-D) Tau aggregates spread and occupy the optic lobe of the brain in 15 days old adult head. (E) Graphical representation of the distance of tau aggregates from the outer boundary of the lamina in 1 and 15 days old flies. ***P<0.001 using student t-test. Scale bar A-D = 50 µm.

## References

1. Vogels T, Leuzy A, Cicognola C, Ashton NJ, Smolek T, Novak M, Blennow K, Zetterberg H, Hromadka T, Zilka N, Schöll M (2019) Propagation of tau pathology: integrating insights from postmortem and in vivo studies. Biol Psychiatry. doi: https://doi.org/10.1016/j.biopsych.2019.09.019.

2. Chang HY, Sang TK, Chiang AS. (2018) Untangling the tauopathy for Alzheimer’s disease and parkinsonism. J Biomed Sci. 25, 54.

3. Orr ME, Sullivan AC, Frost B. (2017) A brief overview of tauopathy: causes, consequences, and therapeutic strategies. Trends Pharmacol Sci. 38, 637–648.

4. Braak H, Braak E. (1991) Neuropathological stageing of Alzheimer-related changes. Acta Neuropathol. 82, 239–259.

5. Albert M, Mairet-Coello G, Danis C, Lieger S, Caillierez R, Carrier S, Skrobala E, Landrieu I, Michel A, Schmitt M, Citron M, Downey P, Courade JP, Buée L, Colin M. (2019) Prevention of tau seeding and propagation by immunotherapy with a central tau epitope antibody. Brain 142, 1736–1750.

6. Takeda S. (2019) Tau propagation as a diagnostic and therapeutic target for dementia: potentials and unanswered questions. Front Neurosci. 13, 1274.

7. Clavaguera F, Bolmont T, Crowther RA, Abramowski D, Frank S, Probst A, Fraser G, Stalder AK, Beibel M, Staufenbiel M, Jucker M, Goedert M, Tolnay M. (2009) Transmission and spreading of tauopathy in transgenic mouse brain. Nat Cell Biol. 11, 909–13.

8. Smolek T, Jadhav S, Brezovakova V, Cubinkova V, Valachova B, Novak P, Zilka N. (2019) First-in-rat study of human Alzheimer’s disease tau propagation. Mol Neurobiol. 56, 621–631.

9. Clavaguera F, Duyckaerts C, & Haïk S. (2020). Prion-like properties of Tau assemblies. Curr Opin Neurobiol. 61, 49–57.

10. Woerman AL, Aoyagi A, Patel S, Kazmi SA, Lobach I, Grinberg LT, McKee AC, Seeley WW, Olson SH, Prusiner SB. (2016) Tau prions from Alzheimer’s disease and chronic traumatic encephalopathy patients propagate in cultured cells. Proc Natl Acad Sci U S A. 113, E8187–E8196.

11. Tanaka Y, Yamada K, Satake K, Nishida I, Heuberger M, Kuwahara T, Iwatsubo T. (2019) Seeding activity-based detection uncovers the different release mechanisms of seed-competent tau versus inert tau via lysosomal exocytosis. Front Neurosci. 13, 1258.

12. Sivanantharajah L, Mudher A, Shepherd D. (2019) An evaluation of *Drosophila* as a model system for studying tauopathies such as Alzheimer’s disease. J Neurosci Methods. 319, 77–88.

13. Iba M, Guo JL, McBride JD, Zhang B, Trojanowski JQ, Lee VM. (2013) Synthetic tau fibrils mediate transmission of neurofibrillary tangles in a transgenic mouse model of Alzheimer’s-like tauopathy. J Neurosci. 33, 1024–37.

14. Wegmann S, Bennett RE, Delorme L, Robbins AB, Hu M, McKenzie D, Kirk MJ, Schiantarelli J, Tunio N, Amaral AC, Fan Z, Nicholls S, Hudry E, Hyman BT. (2019) Experimental evidence for the age dependence of tau protein spread in the brain. Sci Adv. 5, eaaw6404.

15. Wittmann CW, Wszolek MF, Shulman JM, Salvaterra PM, Lewis J, Hutton M, Feany MB. (2001) Tauopathy in *Drosophila*: neurodegeneration without neurofibrillary tangles. Science 293, 711–4.

16. Hay BA, Wolff T, Rubin GM. (1994) Expression of baculovirus P35 prevents cell death in *Drosophila*. Development 120, 2121–9.

17. Hsieh YC, Guo C, Yalamanchili HK, Abreha M, Al-Ouran R, Li Y5, Dammer EB, Lah JJ, Levey AI, Bennett DA, De Jager PL, Seyfried NT, Liu Z, Shulman JM. (2019) Tau-mediated disruption of the spliceosome triggers cryptic RNA splicing and neurodegeneration in Alzheimer’s disease. Cell Rep. 29, 301-316.e10.

18. Freeman M. (1996) Reiterative use of the EGF receptor triggers differentiation of all cell types in the *Drosophila* eye. Cell 87, 651–60.

19. Chanu SI, Sarkar S. (2017) Targeted downregulation of dMyc restricts neurofibrillary tangles mediated pathogenesis of human neuronal tauopathies in *Drosophila*. Biochim Biophys Acta Mol Basis Dis. 1863, 2111–2119.

20. Mencarelli C, Pichaud F. (2015) Orthodenticle is required for the expression of principal recognition molecules that control axon targeting in the *Drosophila* retina. PLoS Genet. 11, e1005303.

21. Santos A, Martín P, Blasco A, Solano J, Cózar B, García D, Goicolea J, Bellas C, Coronado MJ. (2018) NETs detection and quantification in paraffin embedded samples using confocal microscopy. Micron 114, 1–7.

22. Wu TH, Lu YN, Chuang CL, Wu CL, Chiang AS, Krantz DE, Chang HY. (2013) Loss of vesicular dopamine release precedes tauopathy in degenerative dopaminergic neurons in a *Drosophila* model expressing human tau. Acta Neuropathol. 125, 711–25.

23. Greeve I, Kretzschmar D, Tschäpe JA, Beyn A, Brellinger C, Schweizer M, Nitsch RM, Reifegerste R. (2004) Age-dependent neurodegeneration and Alzheimer-amyloid plaque formation in transgenic *Drosophila*. J Neurosci. 24, 3899–906.

24. Babcock DT, Ganetzky B. (2015) Transcellular spreading of huntingtin aggregates in the *Drosophila* brain. Proc Natl Acad Sci U S A. 112, E5427–33.

25. Cleveland DW, Hwo SY, Kirschner MW. (1977) Physical and chemical properties of purified tau factor and the role of tau in microtubule assembly. J Mol Biol. 116, 227–47.

26. Iqbal K, Liu F, Gong CX, Grundke-Iqbal I. (2010) Tau in Alzheimer disease and related tauopathies. Curr Alzheimer Res. 7, 656–664.

27. Kimura T, Sharma G, Ishiguro K, Hisanaga SI. (2018) Phospho-tau bar code: analysis of phosphoisotypes of tau and its application to tauopathy. Front Neurosci. 12, 44.

28. Lindwall G, Cole RD. (1984) Phosphorylation affects the ability of tau protein to promote microtubule assembly. J Biol Chem. 259, 5301–5.

29. Trushina NI, Bakota L, Mulkidjanian AY, Brandt R. (2019) The evolution of tau phosphorylation and interactions. Front Aging Neurosci. 11, 256.

30. Passarella D, Goedert M. (2018) Beta-sheet assembly of tau and neurodegeneration in *Drosophila melanogaster*. Neurobiol Aging. 72, 98–105.

31. Trotter MB, Stephens TD, McGrath JP, Steinhilb ML. (2017) The Drosophila model system to study tau action. Methods Cell Biol. 141, 259–286.

32. Schaeffer V, Lavenir I, Ozcelik S, Tolnay M, Winkler DT, Goedert M. (2012) Stimulation of autophagy reduces neurodegeneration in a mouse model of human tauopathy. Brain 135(Pt 7), 2169–77.

33. Chesser AS, Pritchard SM, Johnson GV. (2013) Tau clearance mechanisms and their possible role in the pathogenesis of Alzheimer disease. Front Neurol. 4, 122.

34. Llorens-Martín M, Jurado J, Hernández F, Avila J. (2014) GSK-3β, a pivotal kinase in Alzheimer disease. Front Mol Neurosci. 7, 46.

35. Jackson GR, Wiedau-Pazos M, Sang TK, Wagle N, Brown CA, Massachi S, Geschwind DH. (2002) Human wild-type tau interacts with wingless pathway components and produces neurofibrillary pathology in *Drosophila*. Neuron. 34, 509–19.

36. Noble W, Planel E, Zehr C, Olm V, Meyerson J, Suleman F, Gaynor K, Wang L, LaFrancois J, Feinstein B, Burns M, Krishnamurthy P, Wen Y, Bhat R, Lewis J, Dickson D, Duff K. (2005) Inhibition of glycogen synthase kinase-3 by lithium correlates with reduced tauopathy and degeneration in vivo. Proc Natl Acad Sci U S A. 102, 6990–5.

37. Hutton M. (2000) Molecular genetics of chromosome 17 tauopathies. Ann N Y Acad Sci. 920, 63–73.

38. Friggi-Grelin F, Coulom H, Meller M, Gomez D, Hirsh J, Birman S. (2003) Targeted gene expression in *Drosophila* dopaminergic cells using regulatory sequences from tyrosine hydroxylase. J Neurobiol. 54, 618–27.

39. Niens J, Reh F, Çoban B, Cichewicz K, Eckardt J, Liu YT, Hirsh J, Riemensperger TD. (2017) Dopamine modulates Serotonin innervation in the *Drosophila* Brain. Front Syst Neurosci. 11, 76.

40. DeVos SL, Corjuc BT, Oakley DH, Nobuhara CK, Bannon RN, Chase A, Commins C, Gonzalez JA, Dooley PM, Frosch MP, Hyman BT. (2018) Synaptic tau seeding precedes tau pathology in human Alzheimer’s disease Brain. Front Neurosci. 12, 267.

41. Frost B, Jacks RL, Diamond MI. (2009) Propagation of tau misfolding from the outside to the inside of a cell. J Biol Chem. 284, 12845–52.

42. de Calignon A, Polydoro M, Suárez-Calvet M, William C, Adamowicz DH, Kopeikina KJ, Pitstick R, Sahara N, Ashe KH, Carlson GA, Spires-Jones TL, Hyman BT. (2012) Propagation of tau pathology in a model of early Alzheimer’s disease. Neuron 73, 685–97.

43. Liu L, Drouet V, Wu JW, Witter MP, Small SA, Clelland C, Duff K. (2012) Trans-synaptic spread of tau pathology in vivo. PLoS One 7, e31302.

44. Mudher A, Shepherd D, Newman TA, Mildren P, Jukes JP, Squire A, Mears A, Drummond JA, Berg S, MacKay D, Asuni AA, Bhat R, Lovestone S. (2004) GSK-3beta inhibition reverses axonal transport defects and behavioural phenotypes in *Drosophila*. Mol Psychiatry 9, 812.

45. Cowan CM, Bossing T, Page A, Shepherd D, Mudher A. (2010) Soluble hyperphosphorylated tau causes microtubule breakdown and functionally compromises normal tau in vivo. Acta Neuropathol. 120, 593–604.

46. Johnson GV, Stoothoff WH. (2004) Tau phosphorylation in neuronal cell function and dysfunction. J Cell Sci. 117, 5721–9.

