## Supplementary figures and images for "Age dependent trans-cellular propagation of human tau aggregates in *Drosophila* disease models"

### Supplemental figure 1

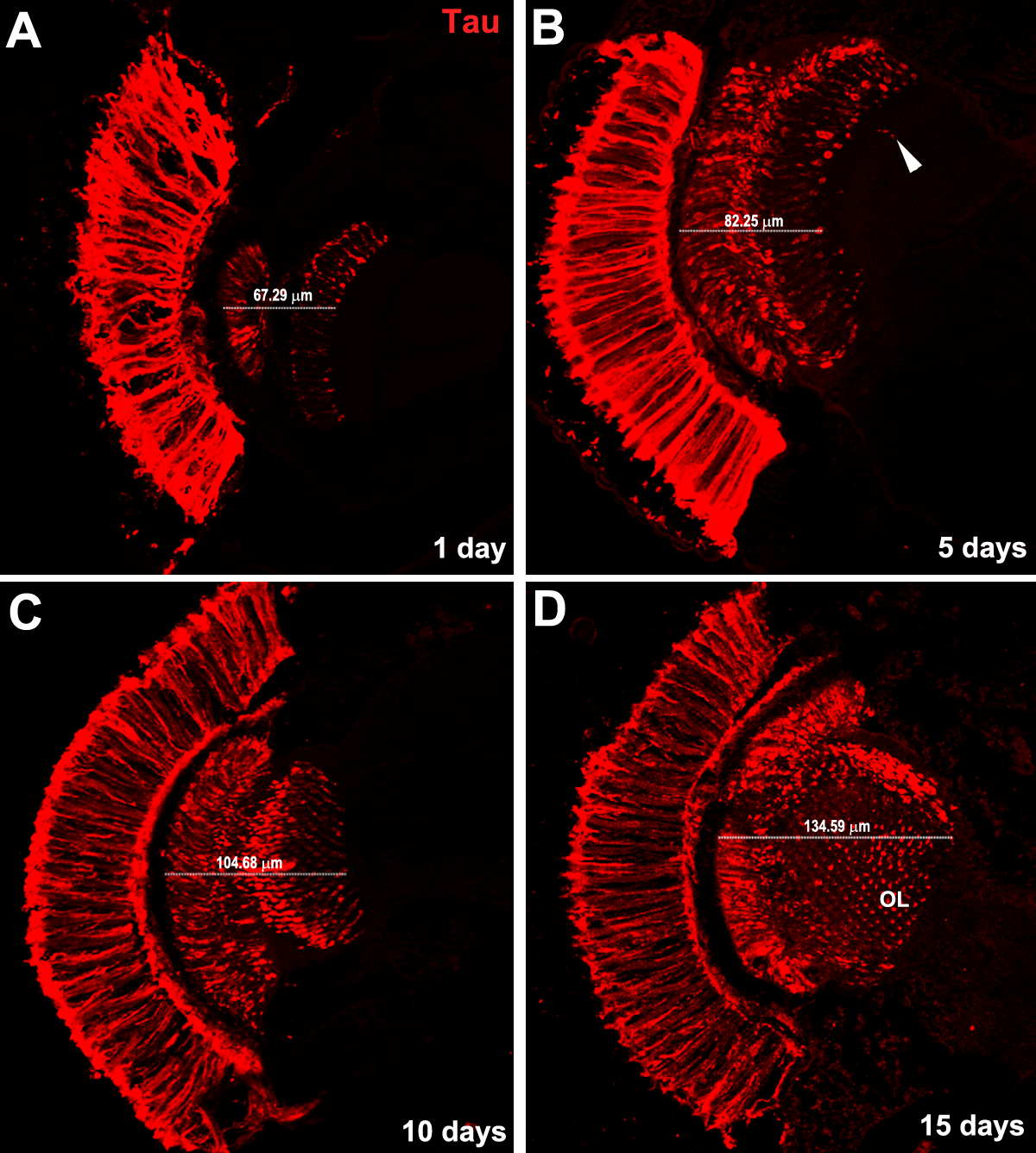

### Supplemental figure 2

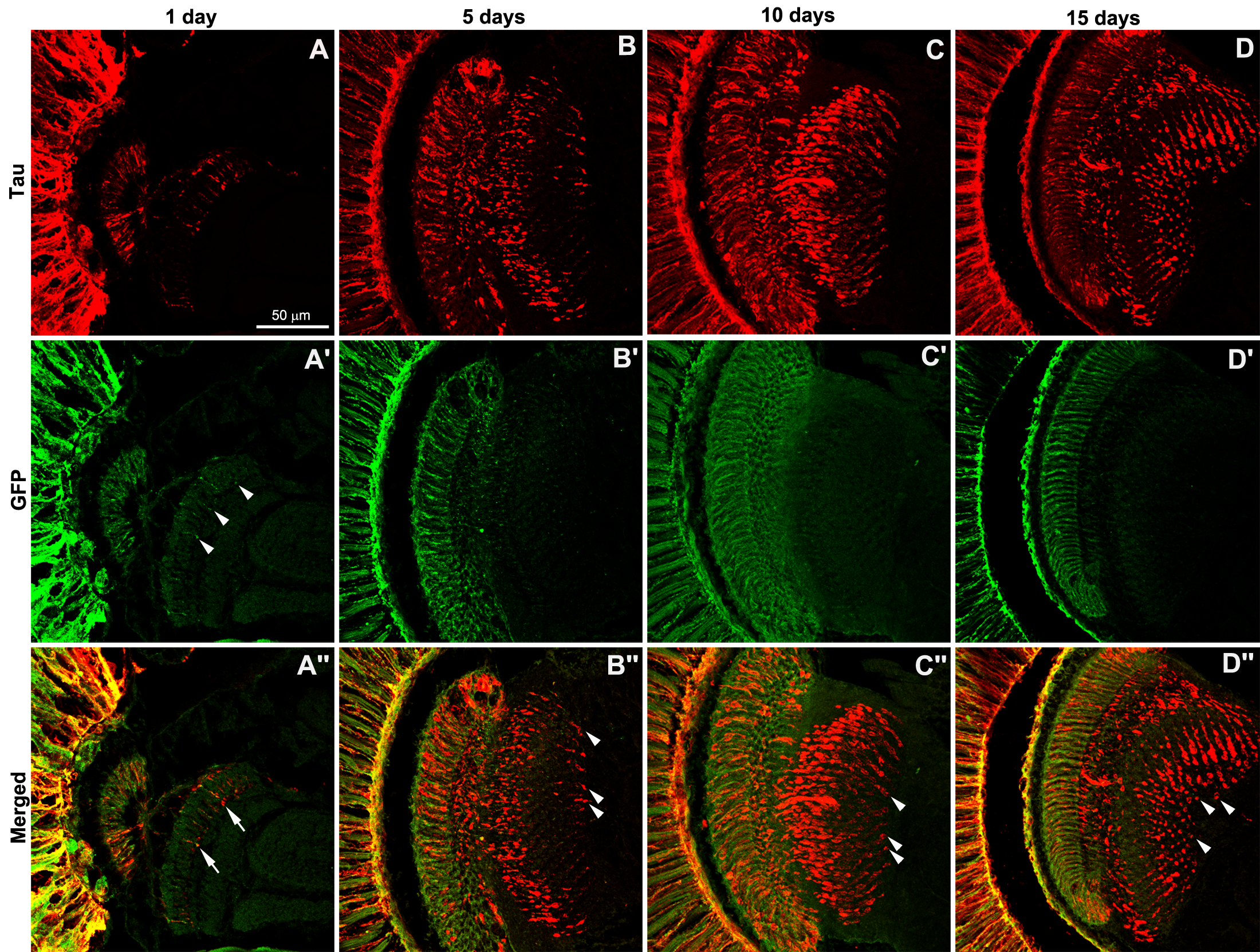

### Supplemental figure 3

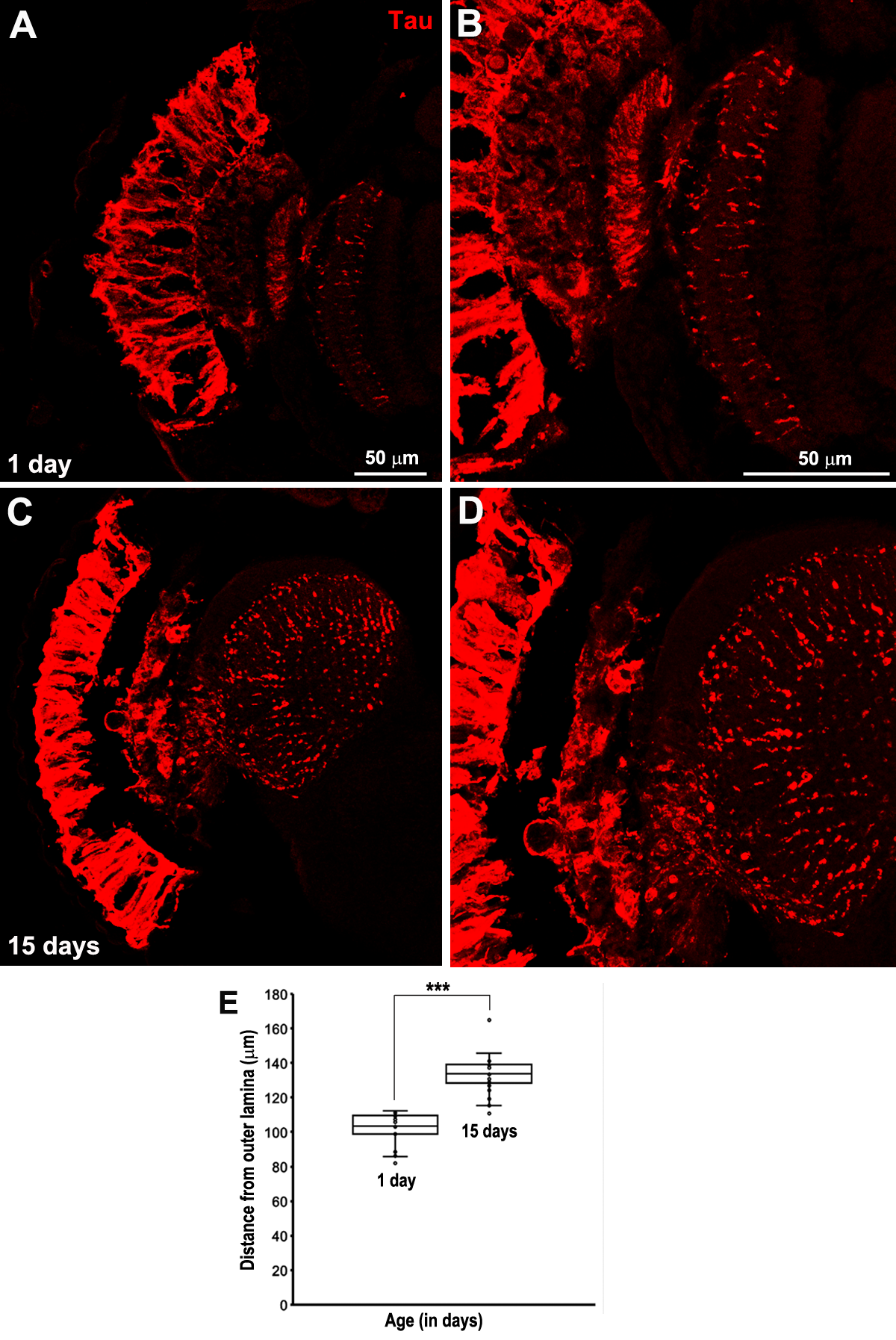
